# Environmental gradients retain the key dimensions of the root economics space at the community level

**DOI:** 10.1101/2025.03.06.641814

**Authors:** Hui Guo, XiaoYu Meng, Yun Zhao, XiaoQing Chen, Lei Wang

## Abstract

The root economics space (RES) provides a framework for understanding species-level belowground strategies, but niche differentiation and environmental filtering may cause community-level root trait coordination to diverge from species-level patterns. Here, we compiled fine-root trait data measured directly from bulk soil samples across 302 plant communities worldwide to examine community-level root trait coordination and its climatic and edaphic drivers. Analyses based on a subset of communities with complete RES traits revealed two major dimensions of variation that were partly consistent with, but not identical to, the classical species-level RES framework. A collaboration-related axis, mainly defined by root diameter and specific root length, was associated with soil pH, with near-neutral soils linked to trait combinations consistent with more “do-it-yourself” nutrient acquisition and acidic soils associated with patterns suggesting greater reliance on mycorrhizal “outsourcing” pathways. A second fast–slow axis, characterized by root tissue density, root nitrogen concentration and root diameter, was associated with precipitation and temperature gradients, with drier environments linked to more conservative trait syndromes and humid, cooler environments associated with more acquisitive strategies. Root nutrient traits also showed ecosystem-specific patterns: in forests, root nitrogen tended to decrease whereas root phosphorus increased along soil nutrient gradients, while in grasslands, root nitrogen increased with soil total nitrogen. Overall, community-level fine-root traits retained key structural elements of the RES framework while exhibiting context-dependent deviations from species-level expectations. These findings highlight how environmental gradients shape belowground trait organization across communities and improve our understanding of plant nutrient-acquisition strategies at large spatial scales.

## Introduction

Functional trait analyses at the community level are pivotal in ecology because they unite species’ traits, abundances, and intraspecific variation to explain how communities assemble and function (Dubuis et al., 2013; Walker et al., 2022). Central to this effort is trait economics—understanding how plants allocate resources among growth, maintenance, and reproduction—since these trade-offs determine ecological strategies across environments (Augusto & Boča, 2022). While species-level frameworks reveal individual plant strategies, ecosystem processes arise from the collective trait composition of coexisting species, rather than those of single taxa (Bardgett et al., 2014; Bruelheide et al., 2018). Therefore, extending trait-economic theory to the community level is critical for uncovering how plant trait coordination responds to environmental gradients and governs ecosystem functioning.

Root traits play a pivotal role in the belowground resource acquisition of plants. The root economics space (RES) framework offers a conceptual model for organizing fine-root traits—including root diameter (RD), specific root length (SRL), root tissue density (RTD), and root nitrogen (RN) concentration—along two main axes: a fast–slow conservation gradient (dominated by RTD and RN) and a collaboration gradient (dominated by RD and SRL) (Bergmann et al., 2020). The RES has been linked to plant resource-use strategies and ecosystem functions such as nutrient cycling and soil carbon dynamics (Freschet et al., 2021). However, niche differentiation fosters complementary resource acquisition among coexisting species (MacArthur & Levins, 1967), potentially causing community-level fine-root trait trade-offs to diverge from species-level patterns (Bruelheide et al., 2018). For example, a field-based study sampling bulk fine roots from 129 local natural communities found that the RD, SRL and RN collapsed into a single major gradient at the community level (Li et al., 2019). Most existing community-level analyses rely on community-weighted means (CWMs) (Matthus et al., 2025). A recent synthesis combining the composition of 810 aboveground species with belowground traits from the Groot database (Guerrero-Ramírez et al., 2021) reported that CWMs align RD, SRL, and RN along a single axis, while RTD loads on a separate axis—deviating from the species-level RES (Barry et al., 2025). Notably, CWM-based approaches typically use fixed species-level trait values and therefore do not capture environment-driven intraspecific variation, potentially obscuring how traits are expressed across environmental gradients.

Community-level root trait–environment relationships are primarily shaped by environmental filtering, which promotes the convergence of trait syndromes under specific environmental conditions (Cadotte et al., 2017), rather than by the microtopographic or microclimatic variation that often drives species-level patterns (Reich et al., 2003). Along global nutrient gradients, the collaboration axis is expected to reflect alternative pathways of nutrient acquisition. In tropical ecosystems, strongly acidic soils generally contain very low levels of plant-available nutrients, particularly phosphorus (Yuan et al., 2011; Du et al., 2020). In response, plants often produce thicker roots that enhance mycorrhizal colonization and strengthen their association with arbuscular mycorrhizal fungi (Kong et al., 2014; Ma et al., 2018). These fungi enhance phosphorus acquisition by extending the effective absorptive surface area beyond the rhizosphere through extensive extraradical hyphal networks (Lambers et al., 2008; Genre et al., 2020). In contrast, temperate forests and grasslands typically experience moderate to higher nutrient availability compared with the severely leached, P-depleted tropical soils (Hinsinger, 2001), plants tend to develop thinner roots with higher specific root length, which allow dense root proliferation and efficient direct nutrient foraging without requiring substantial carbon allocation to mycorrhizal fungi (Chen et al., 2013).

Together, these contrasting environmental filters—reinforced by the geometric linkage between RD and SRL (Zhang et al., 2024)—help maintain a distinct collaboration axis at the community level. The fast–slow conservation axis is likely more complex: RN tends to track soil N availability (Li et al., 2015; Ma et al., 2021; Zhao et al., 2022; Guo et al., 2026), whereas RTD aligns more strongly with climate—especially temperature and precipitation—than with soil nutrients (Freschet et al., 2017; Laughlin et al., 2021). Where these drivers are weakly correlated, the canonical fast–slow axis may fragment at community level.

In this study, we adopted a community-integrated sampling approach, directly measuring fine-root traits from bulk soil samples. This method provides a more realistic representation of belowground structure because CWMs based on species abundances often fail to capture the actual contribution of each species to the fine-root pool—namely, its realized root length or biomass within the soil (Ottaviani et al., 2020; Freschet et al., 2021). Using this approach, we compiled a global dataset encompassing 302 natural plant communities. We applied principal component analysis (PCA) to characterize patterns of trait covariation and identify the dominant dimensions of community-level fine-root economics. Subsequently, we examined how climatic and edaphic factors correspond to these principal axes to uncover the environmental filters that govern belowground strategies at the community level. Specifically, we tested the hypothesis: (a) the community-level fine-root trait coordination diverges from the species-level RES; (b) the collaboration axis, defined by the trade-off between RD and SRL, would persist at the community level and be associated with gradients in soil nutrient conditions; (c) the fast–slow conservation axis, typically represented by RTD and RN, might weaken or fragment, as RTD is more closely regulated by climatic conditions, whereas RN largely responds to soil nutrient status.

## Material and methods

### Data collection

To compile a comprehensive global dataset of community-level fine root traits, we conducted a systematic literature review across three major databases: Web of Science, Google Scholar, and the China National Knowledge Infrastructure (CNKI). The search, covering publications up to October 2025, and the search terms were structured as follows: (“community-level” OR “plant community” OR “forest” OR “grassland” OR “shrubland”) AND (“fine root” OR “root trait” OR “root strategy” OR “nutrient acquisition strategy” OR “root diameter” OR “specific root length” OR “root tissue density” OR “root nitrogen concentration” OR “root phosphorus concentration”).

Studies were included based on the following criteria: (1) Field-based data: Only studies reporting observational data collected in natural plant communities were considered. We excluded studies conducted in greenhouses, pots, hydroponic setups, or heavily disturbed/restored sites. (2) Only studies that explicitly measured fine roots were included. The majority of studies defined fine roots as those with diameters < 2 mm, consistent with common morphological criteria. Studies that instead defined fine roots by developmental order—typically sampling only first-to third-order roots—were also included. No studies using broader or undefined root categories were retained. (3) Trait values had to be reported at the community level, using methods such as bulk sampling through soil cores, root growth cores, or soil pits. Studies presenting species-level values, species-weighted means, or species averages were excluded. (4) Trait completeness: each included study must report at least one of the following traits: RD, SRL, RTD, RN, RP and N:P ratio. This criterion was necessary because morphological traits and chemical traits were often measured in different studies—many reporting only morphological traits and others only chemical traits—so retaining studies with any single trait allowed us to examine how individual traits respond to climate and soil variation. (5) When one of the three morphological traits (RD, SRL, or RTD) was missing, it was estimated using the geometric relationship: SRL=4/(π*RD^2^*RTD). (6) Because fine roots at the community level are predominantly concentrated in shallow soil layers, especially in communities composed of herbaceous and woody plants, where herbaceous roots are mainly distributed in the upper soil, the fine-root trait data collected in this study were restricted to shallow soil layers (≤30 cm). Similarly, soil nutrient variables were also limited to shallow soil layers. (7) Environmental data: Geographic coordinates or location descriptors had to be provided, along with corresponding climate variables (mean annual temperature and precipitation) and soil nutrient properties derived from shallow soil layers, including total N and total P, and soil pH. (8) When multiple local soil measurements were reported under identical climatic conditions, corresponding fine-root trait values were treated as independent observations. Conversely, when a single soil measurement corresponded to multiple fine-root trait measurements, trait values were averaged to match the spatial resolution of soil data. (9) For studies based on experimental designs, only trait values from untreated control plots were used to avoid confounding treatment effects.

Following this selection process, we compiled data from 101 peer-reviewed articles, covering 302 natural plant communities globally. These included forests (n = 202) and grasslands (n = 100). Among these communities, 214 reported at least two fine-root traits, enabling comparative analyses across multiple trait dimensions, and 63 communities provided complete measurements of RD, SRL, RTD, and RN. The distribution of sampling sites across biomes is illustrated in Fig. S1.

### Multivariate analysis of trait coordination

To meet assumptions of normality and homoscedasticity, all fine-root trait data were log_10_-transformed prior to analysis. Of the compiled dataset, 63 communities contained complete measurements of the four key RES traits—RD, SRL, RTD, and RN—and were therefore used for PCA to identify the major axes of trait coordination. We additionally examined pairwise correlations among these four traits within the 63-community subset and compared them with correlations from the full dataset to evaluate the robustness and generality of observed trait relationships. Finally, we related the first and second principal component (PC) axes to climatic and edaphic variables using regression models to assess how environmental filtering shapes community-level fine-root strategies.

## Results

### Community-level root trait relationships and root economic axes

The correlations between root morphological and chemical traits were weak. Correlation analyses revealed a strong negative relationship between RD and SRL (*r* = −0.74, *p* < 0.001; Fig. 1a). Both RD and SRL were also significantly negatively correlated with RTD (*r* = −0.24, *p* < 0.01; *r* = −0.23, *p* < 0.01, respectively). In contrast, RTD showed no significant correlation with RN (*r* = −0.16, *p* = 0.21; Fig. 1a). Among chemical traits, RN showed a weak positive association with RP (*r* = 0.28, *p* = 0.06; Fig. 1a). Because the N:P ratio is derived directly from nitrogen and phosphorus concentrations, its positive correlation with RN (*r* = 0.57, *p* < 0.001) and negative correlation with RP (*r* = −0.71, *p* < 0.001) are expected.

**Figure 1.**
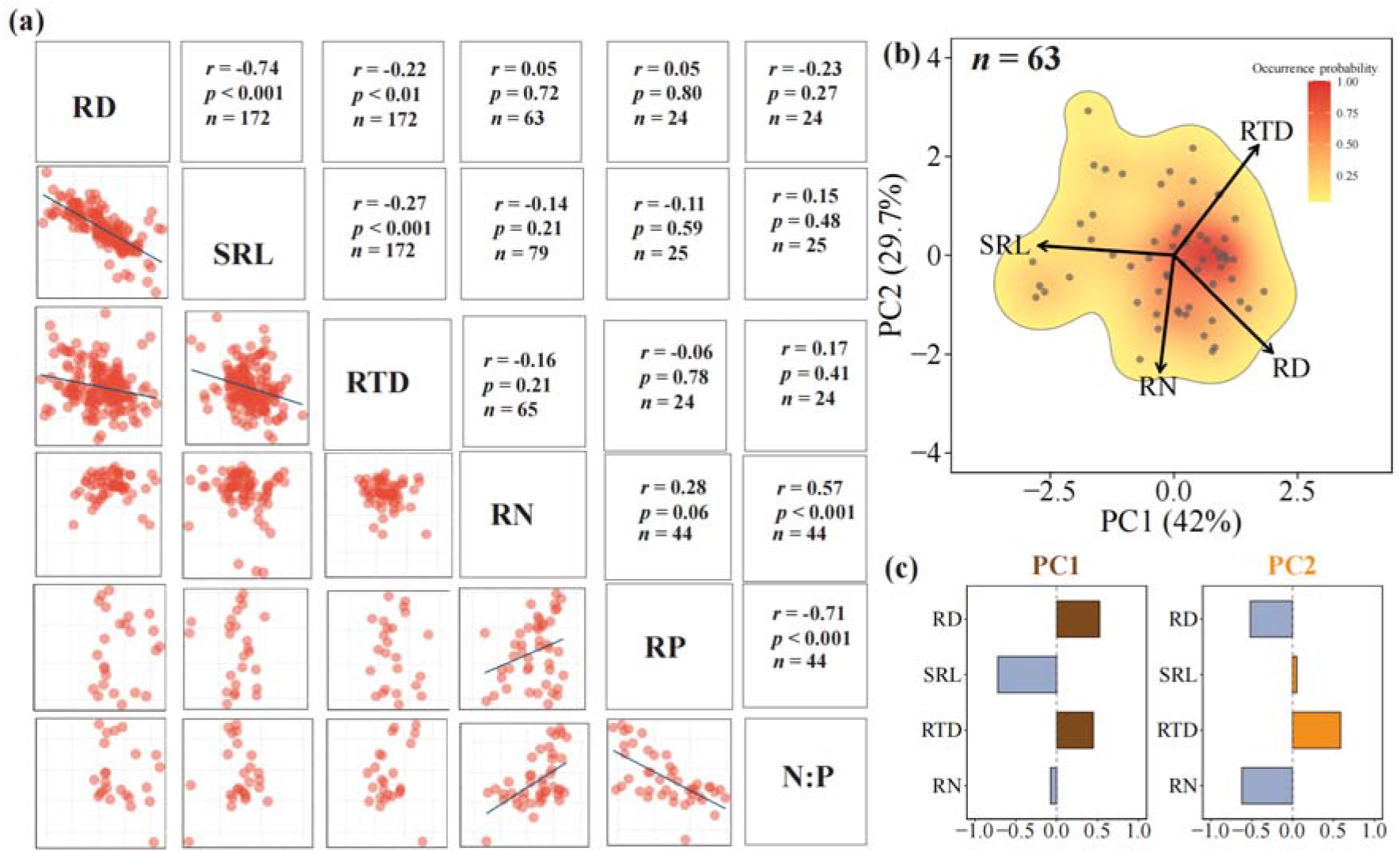
Community-level coordination of fine-root traits and principal component structure of the root economics space. (a) Pairwise Pearson’s correlations among community-level fine-root traits. The lower panels show bivariate relationships between traits, and the upper panels show correlation coefficients, significance levels, and sample sizes. Morphological traits were strongly coordinated, particularly the negative relationship between root diameter and specific root length, whereas correlations between morphological and nutrient traits were generally weak. (b) Principal component analysis of communities with complete measurements of root diameter, specific root length, root tissue density, and root nitrogen concentration (n = 63). Colour gradients indicate the probability density of plant communities within the PCA-defined root trait space, with red indicating high-density regions and yellow indicating low-density regions. PC1 was mainly defined by the RD–SRL trade-off, representing the collaboration axis, whereas PC2 was primarily associated with RTD, RN, and RD, representing the fast–slow axis. (c) Trait loadings on PC1 and PC2, showing the relative contribution and direction of each trait along the two principal components. Trait abbreviations: RD, root diameter; SRL, specific root length; RTD, root tissue density; RN, root nitrogen concentration; RP, root phosphorus concentration; N, root nitrogen-to-phosphorus ratio.

PCA of the 63 communities showed that community-level root trait organization retained the key dimensions of the species-level root economics space (RES). The first principal component (PC1) was primarily defined by RD and SRL, representing the collaboration axis, whereas the second component (PC2) retained the fast–slow gradient dominated by RTD and RN, with RD also contributing substantially (Fig. 1b, c; Table S1). Correlation analyses within this 63-site subset showed patterns consistent with those from the full dataset: RN was not significantly correlated with RD, SRL, or RTD, and RD and RTD were likewise uncorrelated (Fig. S2), supporting the robustness of the identified trait relationships.

### Effects of climate and soil factors on root economic axes

PC1, representing the community-level collaboration axis, was most strongly associated with soil pH (*r*^2^ = 0.31, *p* < 0.001; Fig. 2a). This relationship indicates that variation in soil chemical conditions was linked to shifts from trait combinations associated with mycorrhizal outsourcing in more acidic soils toward more root-based “do-it-yourself” nutrient acquisition in near-neutral soils. Mean annual temperature was also significantly related to PC1 (*r*^2^ = 0.15, *p* < 0.01; Fig. 2b), suggesting that temperature acted as an additional filter on the collaboration axis. In contrast, the PC2, representing the fast–slow axis, was mainly associated with mean annual precipitation (*r*^2^ = 0.34, p < 0.001; Fig. 2c), with temperature also exerting a significant effect (*r*^2^ = 0.13, *p* < 0.01; Fig. 2d).

**Figure 2.**
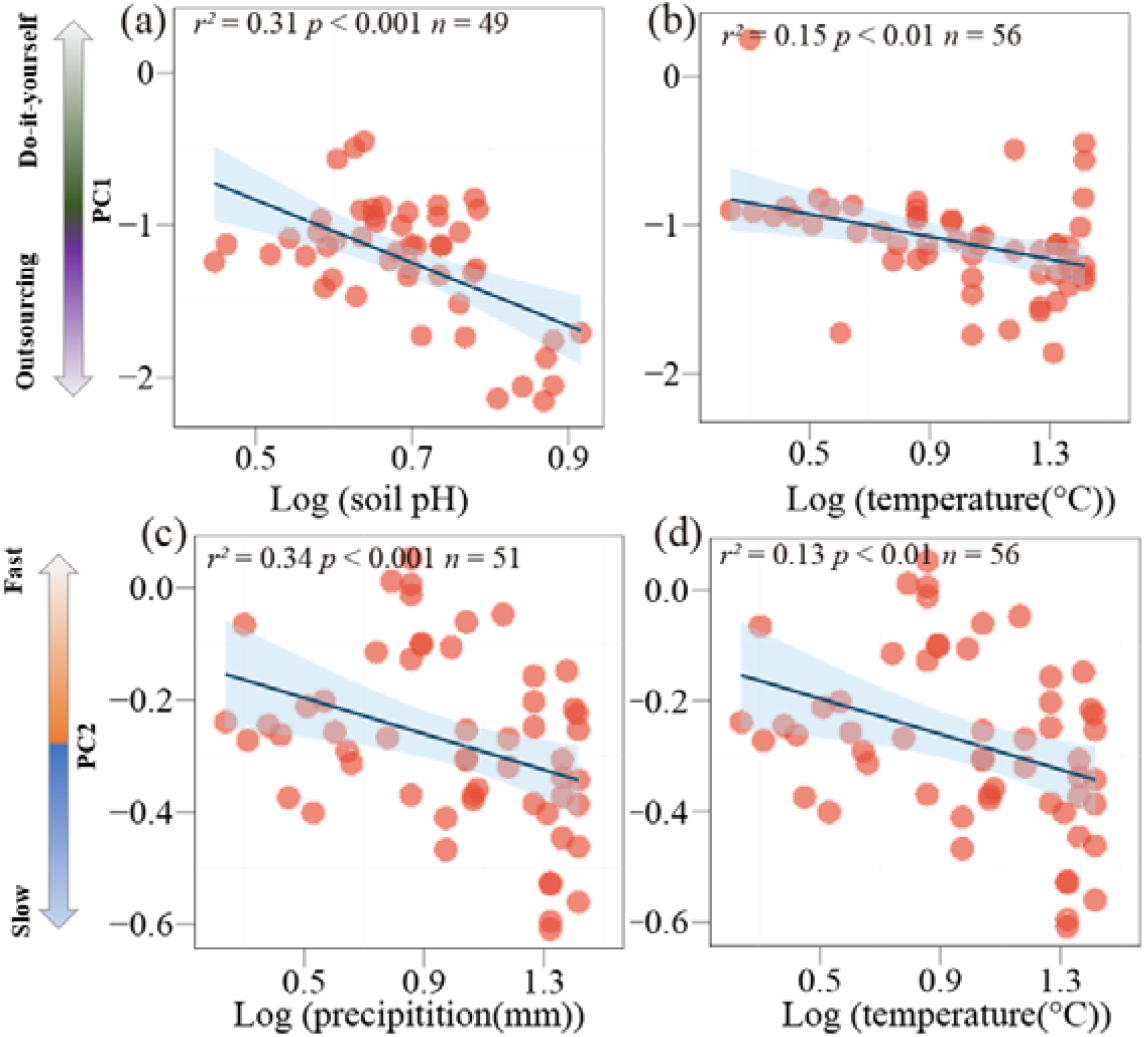
Environmental controls on community-level root economic axes. Relationships between the first two principal components of community-level fine-root traits and major environmental gradients. (a, b) The first principal component (PC1), representing the collaboration axis, was significantly associated with soil pH and temperature. Along this axis, communities shifted from trait combinations associated with greater reliance on mycorrhizal outsourcing under more acidic soil conditions toward more root-based “do-it-yourself” nutrient acquisition under near-neutral soil conditions. Temperature also contributed to variation along this collaboration axis. (c, d) The second principal component (PC2), representing the fast–slow axis, was mainly associated with precipitation and temperature. Drier and warmer environments were linked to more conservative root trait syndromes, whereas wetter and cooler environments were associated with more acquisitive strategies. Blue lines indicate fitted linear regressions, and shaded areas represent 95% confidence intervals. All variables were log10-transformed before analysis.

In forest communities, root morphological traits were strongly structured by soil nutrient availability: RD decreased significantly with increasing total soil phosphorus (*r*^2^ = 0.19, *p* < 0.05; Fig. S3a), whereas SRL increased with higher total soil nitrogen (*r*^2^ = 0.14, *p* < 0.01; Fig. S3b). In contrast, RTD declined with increasing precipitation (*r*^2^ = 0.06, *p* < 0.05; Fig. S3c).

### Divergent root N and P responses reveal distinct nutrient-use strategies between forests and grasslands

As soil nutrient availability increased, root nutrient concentration responses exhibited contrasting patterns both between nutrients and across ecosystems. In forest communities, RN declined with increasing soil total nitrogen and soil pH (*r*^2^ = 0.19, *p* < 0.001, Fig. 3a; *r*^2^ = 0.25, *p* < 0.01, Fig. S4a), whereas RP rose markedly with higher levels of both soil total nitrogen (*r*^2^ = 0.40, *p* < 0.01; Fig. 3c) and total phosphorus (*r*^2^ = 0.31, *p* < 0.01; Fig. 3d). Consistently, RN in forest communities was negatively correlated with both soil nitrogen and soil phosphorus availability (*r*^2^ = 0.49, *p* < 0.001; Fig. S5a; *r*^2^ = 0.38, *p* < 0.01; Fig. S5b). In contrast, grassland roots showed a positive relationship between RN and total soil nitrogen and soil pH (*r*^2^ = 0.25, *p* < 0.05; Fig. 3b; *r*^2^ = 0.31, *p* = 0.06; Fig. S4b).

**Figure 3.**
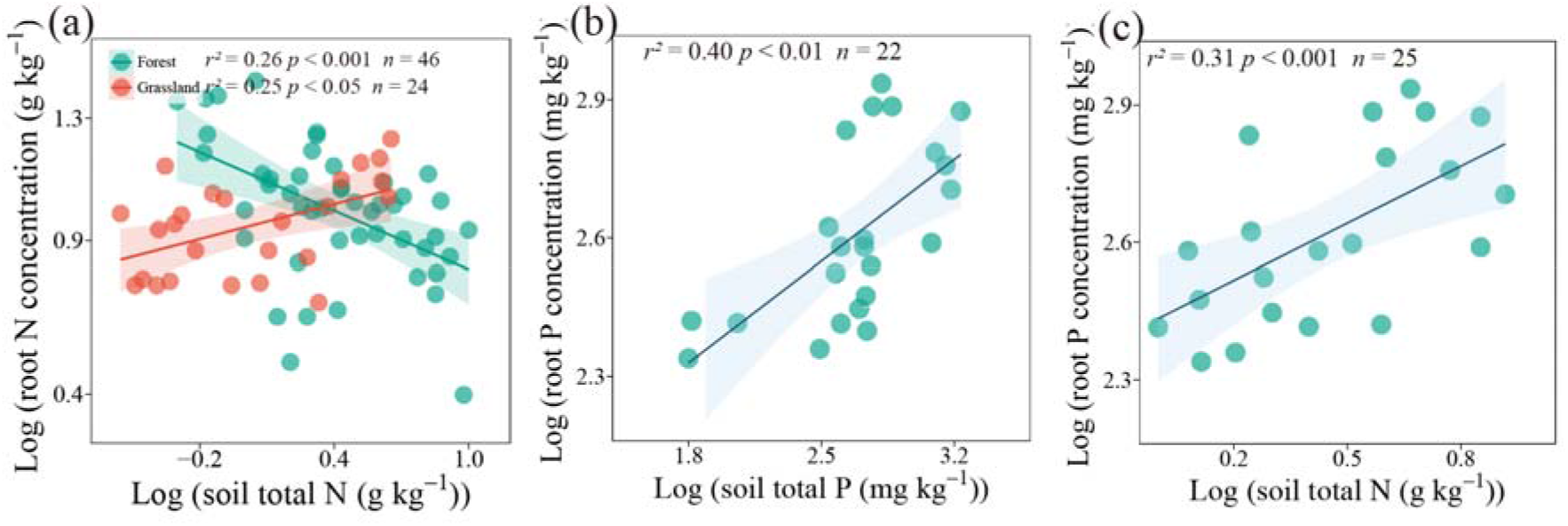
Ecosystem-specific responses of fine-root nutrient concentrations to soil nutrient availability. (a) Relationships between fine-root nitrogen concentration and soil total nitrogen in forest and grassland communities. Forest communities showed a significant negative relationship between root nitrogen concentration and soil total nitrogen, whereas grassland communities showed a significant positive relationship, indicating contrasting nitrogen-use strategies between the two ecosystem types. (b, c) Relationships between fine-root phosphorus concentration and soil total phosphorus and total nitrogen in forest communities. Root phosphorus concentration increased significantly with both soil total phosphorus and soil total nitrogen, suggesting that forest communities may enhance phosphorus accumulation in root tissues under nutrient-enriched conditions. Lines indicate fitted linear regressions, and shaded areas represent 95% confidence intervals. All variables were log10-transformed before analysis.

## Discussion

### Community-level root economics preserves key RES dimensions

Contrary to our initial hypothesis (a), although community-level fine-root economics showed some deviations from species-level patterns, PCA of the 63 communities recovered both a collaboration axis and a fast–slow axis, indicating that community-level fine-root trait organization can retain the core structural dimensions of the species-level RES framework (Bergmann et al., 2020). This result is consistent with a recent study based on artificially assembled communities, which also identified a recognizable RES using bulk-soil trait measurements (Hennecke et al., 2025). The collaboration axis was primarily defined by RD and SRL, which exhibited a strong negative relationship and high absolute loadings in the PCA, indicating that the RD–SRL trade-off remains a robust organizing principle even at the community level.

Importantly, our analyses also recovered a fast–slow axis dominated by RTD and RN, although RD additionally contributed to this dimension. Rather than contradicting the RES framework, this pattern suggests that at the community level RD may participate in both collaboration and resource-conservation dimensions, reflecting a modest structural modification rather than a breakdown of the RES. Notably, several previous studies have failed to detect a clear fast–slow axis at the community level or have reported configurations in which RN aligns with morphological traits such as RD and SRL (Prieto et al., 2016; Li et al., 2019; Borden et al., 2021; Yang et al., 2024; Barry et al., 2025). In contrast, our results show that the canonical fast–slow dimension can still emerge when community-level traits are derived from bulk-soil measurements, highlighting that the absence of this axis is not universal. Although RTD and RN co-loaded on the second axis, their pairwise correlation was weak, suggesting that trait coordination along the fast–slow dimension may be less tightly constrained at the community level than at the species level. Such loosened coordination likely reflects stronger influences of environmental heterogeneity and species composition, which may partly explain discrepancies among previous community-level studies (Matthus et al., 2025). Our findings also differ from global CWM-based syntheses that reported a decoupling of RN and RTD (Barry et al., 2025). This contrast probably arises from differences in ecological representation and methodology: our dataset is dominated by forest communities, many from China, whereas CWM-based syntheses include a larger proportion of grasslands and nitrogen-fixing species, and rely on abundance-weighted traits rather than directly measured bulk-soil root pools. Together, these results suggest that community-level root economics retains the fundamental RES structure, but that the expression and coupling strength of trait axes are context-dependent and sensitive to sampling and analytical approaches.

### Environmental controls on collaboration and fast–slow axes at the community level

Partly consistent with hypothesis (b), PC1—dominated by RD and SRL—was associated swith variation in soil pH (Fig. 2a), suggesting that the collaboration axis may be related to environmental differences linked to soil chemical conditions. Soil pH is known to influence nutrient solubility (Hinsinger, 2001; Marschner, 2012), and the sites included in our dataset covered a range of mostly acidic to near-neutral soils (pH 2.8–7.6), with only one weakly alkaline site (pH = 8.2). In acidic soils (pH < 6.0), elevated concentrations of Fe^3+^ and Al^3+^ promote the formation of insoluble Fe–P and Al–P complexes, strongly limiting phosphorus availability and, together with intense leaching, resulting in generally low levels of plant-available nutrients (Hinsinger, 2001). Under such nutrient-poor conditions, plants may develop thicker roots that provide greater cortical space for arbuscular mycorrhizal fungal colonization (Kong et al., 2014; Ma et al., 2018). By deploying a dense network of extraradical hyphae, these fungi exploit microsites rich in mineral nutrients beyond the rhizosphere and facilitate their subsequent transfer to plant roots (Lambers et al., 2008; Genre et al., 2020). In contrast, near-neutral soils (6.5 < pH ≤ 7.6) typically exhibit higher availability of inorganic nitrogen and phosphorus (Hinsinger, 2001). In these environments, plants can efficiently acquire nutrients through direct root uptake, reducing reliance on mycorrhizal partners and the need for substantial carbon investment in symbiosis (Chen et al., 2013; Fellbaum et al., 2014; Kokkoris, 2025). Consistent with this interpretation, we found that RD in forest communities decreased with increasing soil phosphorus (Fig. S3a), whereas SRL increased with rising soil nitrogen (Fig. S3b), indicating a shift toward a more “do-it-yourself” nutrient acquisition strategy as soil fertility improves. This pattern aligns well with findings from 129 natural communities in southwest China, where soil nutrient gradients strongly shaped RD and SRL (Li et al., 2019). Although this range captures substantial variation in soil conditions, the limited representation of alkaline sites suggests that the inferred relationships between soil pH and root trait expression should be interpreted cautiously. Nevertheless, our results contrast with pot-based experiments using CWMs, which reported that plants in acidic soils rely more on direct root uptake than on mycorrhizal fungi (Lachaise et al., 2022). This discrepancy may arise because, in pot systems, rapid shifts in nitrate-to-ammonium ratios can strongly and artificially alter soil pH (Dotaniya et al., 2015), thereby influencing short-term mycorrhizal responses. In addition, a CWMs-based field study had reported no significant effect of soil pH on the collaboration axis, instead identifying topographic factors as the dominant drivers (Da et al., 2023), this discrepancy may reflect scale-dependent environmental filtering (Cadotte et al., 2017), whereby the influence of soil pH is more apparent across broad gradients but less detectable at local scales.

Temperature acts as an additional filter on the collaboration axis. In warm climates, plant communities tend to develop thicker, lower-SRL roots, indicating a greater reliance on arbuscular mycorrhizal fungi. High temperatures accelerate organic matter decomposition and nutrient turnover (Conant et al., 2011), creating conditions in which mycorrhizal fungi—via their extensive extraradical hyphae—can intercept and acquire inorganic nutrients more efficiently than roots alone (Lambers et al., 2008; Steidinger et al., 2019; Genre et al., 2020), thereby increasing plant dependence on fungal-mediated uptake. In contrast, colder climates favor thin, high-SRL roots, reflecting a shift toward “do-it-yourself” nutrient acquisition. Low temperatures limit photosynthetic carbon supply (Sage et al., 2007), and plant nutrient demand is generally lower than in warm environments, while the relative carbon cost of maintaining mycorrhizal symbiosis remains high (Fellbaum et al., 2014; Kokkoris, 2025). Under such constraints, plants tend to construct low-cost roots to conserve carbon and expand soil exploration, and simultaneously reduce their investment in mycorrhizal partners, thereby lowering the energetic burden of symbiosis.

Contrary to our hypothesis (c), the fast–slow axis was retained at the community level. This axis, represented by PC2, was primarily defined by RTD, RN, and RD and was jointly driven by two climatic factors—precipitation and temperature (Fig. 2c, d). In arid environments characterized by low precipitation and high temperatures, communities tend to exhibit a “conservative” root trait syndrome, including thinner roots, higher RTD, and lower RN. This syndrome likely arises because drought constrains plant productivity (Farooq et al., 2012), thereby favoring more economical, carbon-saving resource acquisition strategies. Thinner roots reduce construction costs by adopting smaller diameters, and higher RTD and lower RN further extend root lifespan and reduce metabolic activity (Bergmann et al., 2020; Huang et al., 2024; Hou et al, 2024), enabling plants to maintain function by low carbon invest under prolonged water limitation. Conversely, in humid, cooler environments, communities more often display an “acquisitive” syndrome with thicker roots, low RTD, and high RN. Abundant moisture and lower temperatures slow microbial activity and nutrient mineralization (Conant et al., 2011; Steidinger et al., 2019), resulting in seasonally pulsed nutrient supply. Plants counter this with high metabolic capacity—manifested as low RTD and high RN—to rapidly exploit nutrients during brief favorable periods. And thicker roots, associated with wider vascular conduits, further enhance hydraulic conductivity and nutrient transport efficiency (Kong et al., 2014; Kong et al., 2021). In contrast to our findings, a CWMs-based study along an elevational gradient reported that the fast–slow axis was mainly driven by soil factors (Da et al., 2023), suggesting that the relative importance of climatic versus edaphic controls may vary across environmental contexts.

### Ecosystem-specific nitrogen responses and stoichiometric regulation of fine roots

Interestingly, soil nitrogen availability elicits contrasting responses across ecosystems (Fig. 3a, b). Unlike species-level findings which reported increased root N with higher soil nitrogen (Nadelhoffer et al., 1985; Ding et al., 2020; Yan et al., 2022), in forests communities, RN decreases with increasing soil nitrogen, likely due to a nitrogen-dilution effect (Jarrell et al., 1981), where nitrogen inputs stimulate biomass growth that outpaces nitrogen accumulation, reducing RN. Similar dilution effects have been reported for leaf and root tissues under nitrogen enrichment (Güsewell, 2004; Lu et al., 2011). Besides, forest plants may also allocate more nitrogen to high-metabolic tissues like leaves or mycorrhizal fungal to balance photosynthesis and respiration (Valverde-Barrantes et al., 2017; Phillips et al., 2013). In grasslands, RN increases with rising soil nitrogen, as short-lived, fast-turnover roots respond rapidly to nutrient enrichment by elevating RN and metabolic activity (Bai et al., 2010; Wang et al., 2019), reflecting a typically acquisitive strategy. These differences explain opposing RN trends with pH-driven nutrient enrichment (Fig. S4) and decreasing forest N:P ratios with rising nitrogen (Fig. S5a). RP significantly increases with rising soil nitrogen and phosphorus levels (Fig. 3b, c), suggesting that in nitrogen-rich environments, forest plants enhance phosphorus uptake to alleviate potential phosphorus limitation (Aerts & Chapin, 2000; Deng et al., 2017; Yuan & Chen, 2015), thereby maintaining nitrogen–phosphorus stoichiometric balance.

Notably, RN exhibits strong positive correlations with RP and the N:P ratio, but weaker associations with morphological traits (RD, SRL, RTD) (Fig. 1a). This decoupling from morphology, together with the tight coupling between N- and P-related traits, suggests that root nutrient concentrations are regulated to maintain internal stoichiometric balance rather than passively tracking structural variation. Specifically, nitrogen supports protein synthesis and metabolic activity, whereas phosphorus underpins energy transfer and nucleic acid metabolism (Yuan et al., 2011; Du et al., 2020). The synchronized variation of RN and RP therefore reflects plants’ capacity to actively coordinate N and P uptake and allocation to maintain optimal N:P ratios, balancing metabolic demand and physiological homeostasis across environmental gradients (Weemstra et al., 2016). This finding underscores that root nutrient concentrations not only align with the RES fast–slow axis but also highlight plants’ active regulation of nitrogen and phosphorus uptake and allocation under diverse environmental constraints.

## Conclusion

Based on a globally compiled dataset, this study adds new insights into community-level root trait organization, showing that it retains key structural elements of the species-level RES while also displaying context-dependent deviations. The collaboration axis defined by RD and SRL remained evident and was associated with variation in soil pH, suggesting that differences in soil chemical environments may influence shifts in nutrient-acquisition strategies across communities. A fast–slow axis involving RTD, RN, and RD was also identified and was associated with climatic gradients, particularly precipitation and temperature, linking drier environments to more conservative trait syndromes and humid, cooler environments to more acquisitive strategies. In addition, forests and grasslands showed contrasting nutrient-related patterns, with forest RN tending to decline while grassland RN increased along soil nitrogen gradients. Overall, Together, these findings indicate that large-scale environmental gradients organize community-level belowground trait structure in ways that both preserve and modify the classical RES, with implications for predicting nutrient-acquisition strategies and ecosystem functioning under environmental change.

## Acknowledgements

This study was supported by the Natural Science Foundation of Xiamen, China (grant nos. 3502Z202471075), the Fujian Provincial Department of Education Project for Young and Middle-aged Teachers (Grant nos. JZ240064), and the Xiamen University of Technology High Level Talent Project (grant nos. YKJ24012R).

## Author contributions

H.G. conceived the idea, performed the data analysis, and wrote the manuscript. X.Y.M, Y.Z., and X.Q.C, L.W. substantially contributed to revisions.

## Data Availability Statemen

All data supporting the findings of this study can be accessed in Dryad database (http://datadryad.org/share/LINK_NOT_FOR_PUBLICATION/OeDbrJ34yd3_hQyNnXDXh1P1q1Va-0ntEffki_bJF0Q).

## Supplementary materials

**Figure S1.**
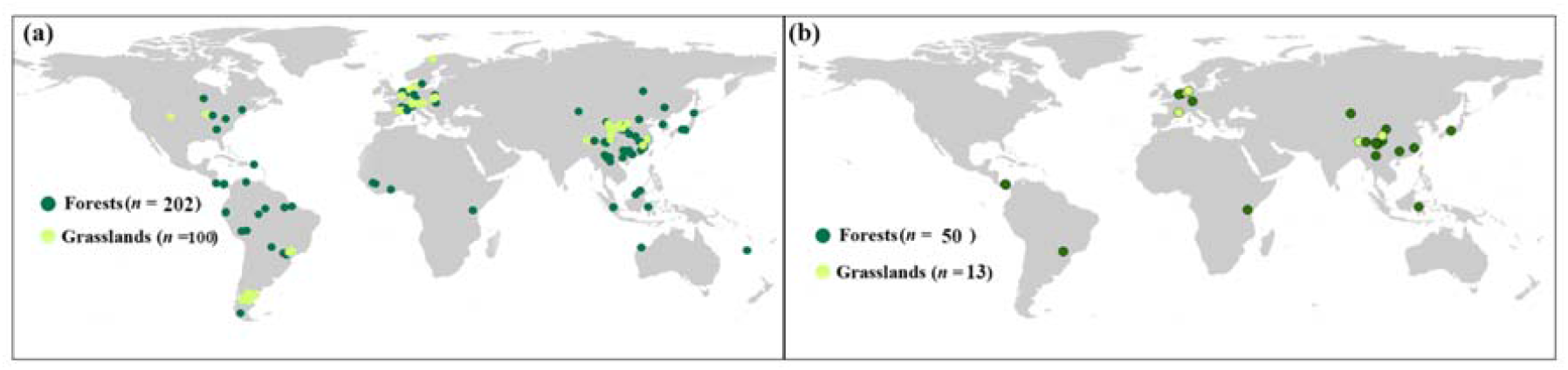
Global Locations of sampling sites. (a) Locations of 302 community-level sampling sites for root traits across forests and grasslands. (b) Locations of 63 community-level sites from forests and grasslands with complete measurements of root diameter, specific root length, root tissue density, and root nitrogen concentration.

**Figure S2.**
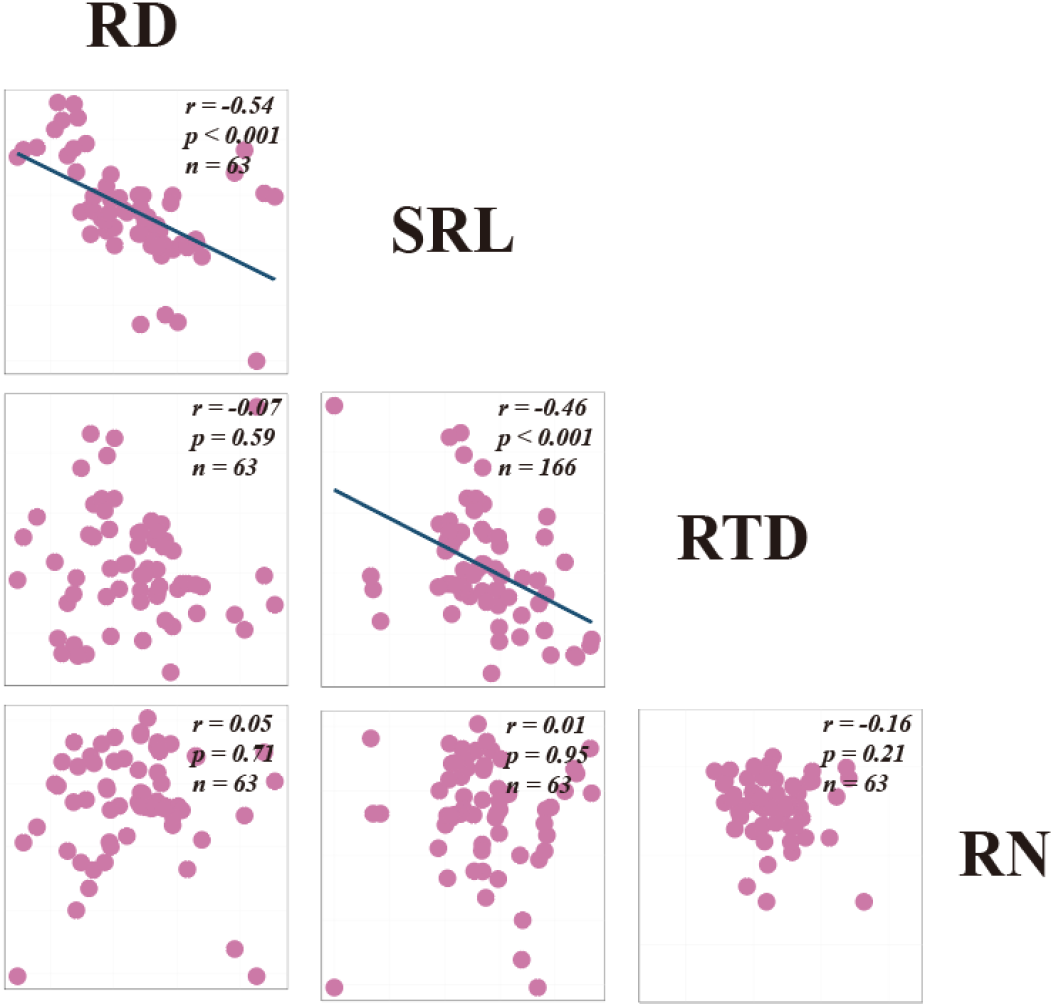
Correlations among fine-root traits in communities with complete RES data (n = 63). The results show that, except for the non-significant relationship between root diameter (RD) and root tissue density (RTD), the correlation patterns among fine-root traits were largely consistent with those derived from all 302 communities, indicating strong robustness and consistency of the dataset. All variables were log10-transformed; Trait abbreviations: SRL, specific root length; RN, root nitrogen concentration.

**Figure S3.**
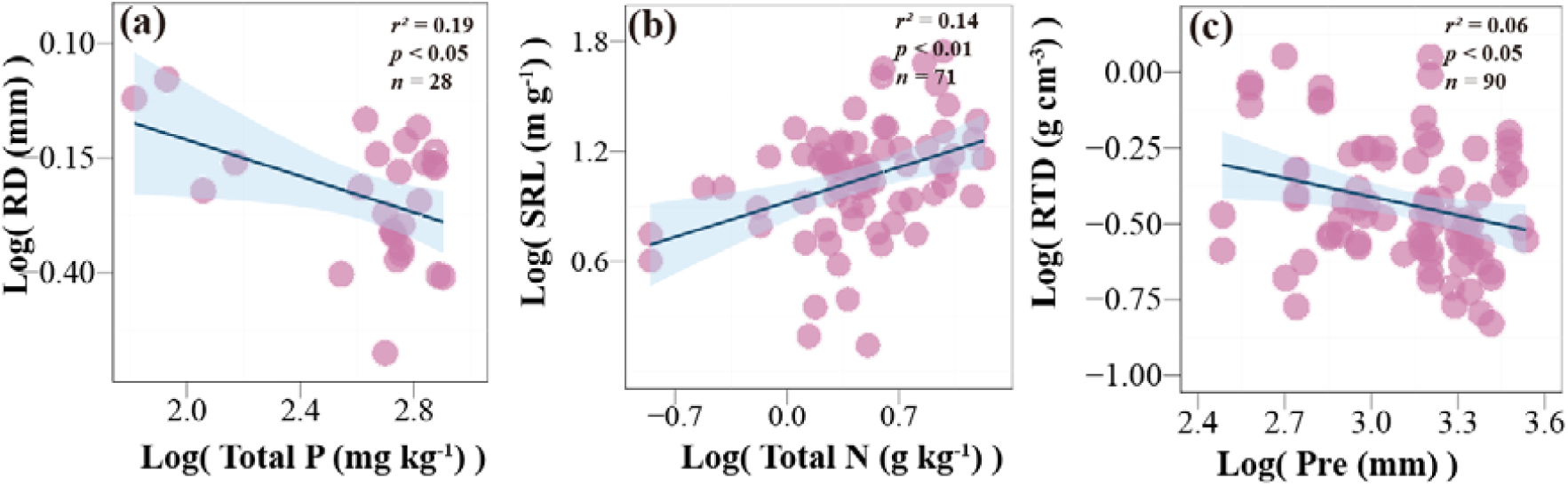
Effects of environmental variables on root diameter (RD, a), specific root length (SRL, b), and root tissue density (RTD, c) in forest communities. The collaboration axis formed by RD and SRL was primarily regulated by soil nutrient status: in nutrient-poor soils, forest plants relied more on mycorrhizal fungi for nutrient mobilization and uptake, whereas in nutrient-rich conditions, they favored direct nutrient absorption through roots—indicating a strategic shift from symbiotic to self-sufficient nutrient acquisition.

**Figure S4.**
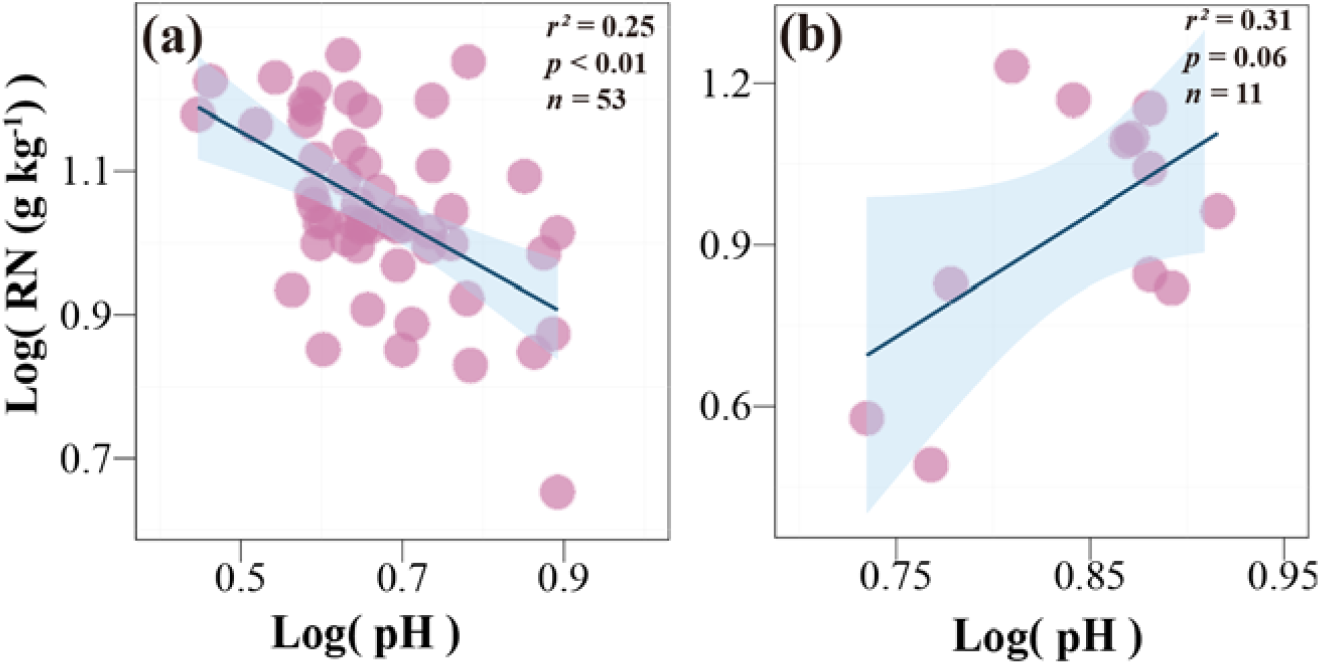
Soil pH induces opposite trends in fine-root nitrogen concentration (RN) between forest (a) and grassland (b) communities. This pattern indicates that in forests, higher soil pH alters nutrient availability and triggers a nitrogen dilution effect—where increased nutrient supply leads to reduced root nitrogen concentration. In contrast, grassland communities do not exhibit this effect; instead, fine-root nitrogen tends to increase with higher soil nutrient availability.

**Figure S5.**
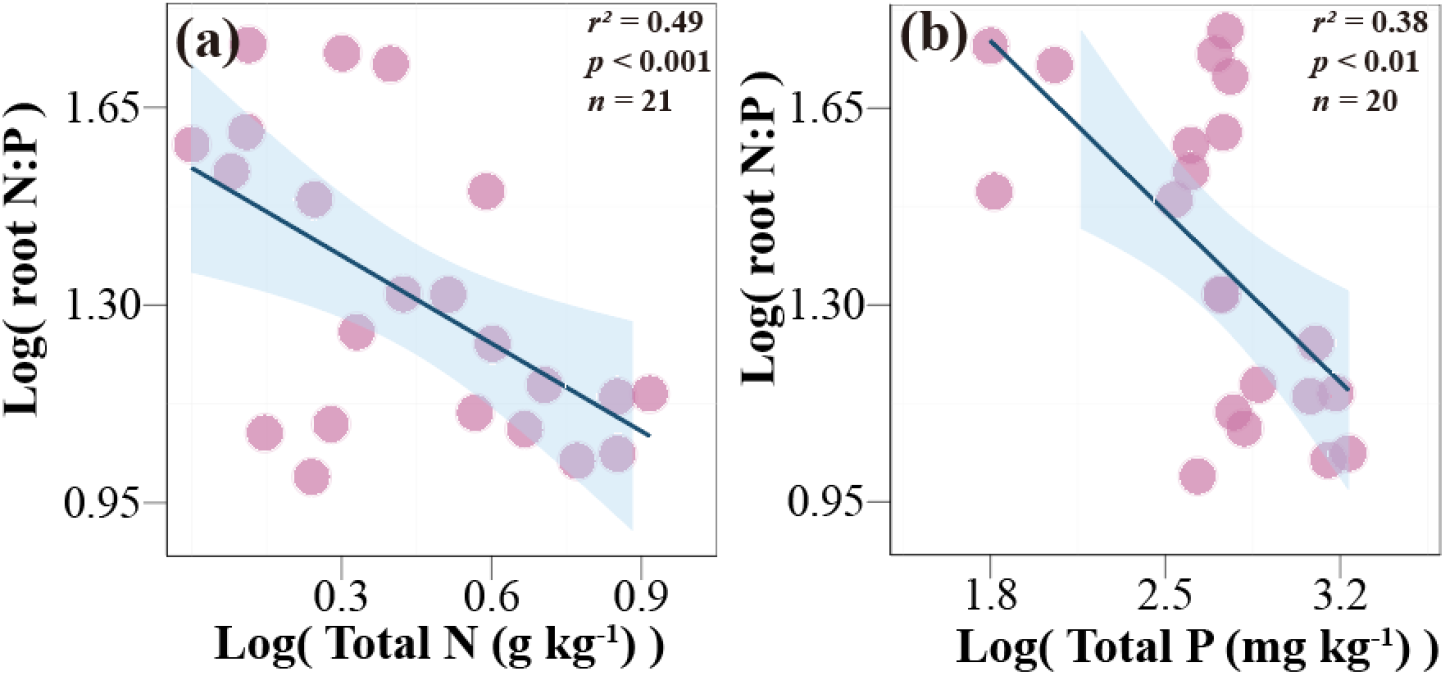
Relationships between fine-root N:P ratio and soil total nitrogen (a) and total phosphorus (b) in forest communities. The fine-root N:P ratio decreased with increasing soil total nitrogen, consistent with a nitrogen dilution effect, whereby nitrogen enrichment enhances root biomass production but dilutes nitrogen concentration, leading to a lower N:P ratio.

**Table S1.** Eigenvalues, proportion of variance and loadings of traits on the principal components (PC1 and PC2).

|  | **PC1** | **PC2** |
| --- | --- | --- |
| % Variance | 42.0 | 29.7 |
| RD | 0.524 | -0.519 |
| SRL | -0.720 | 0.051 |
| RTD | 0.448 | 0.586 |
| RN | -0.075 | -0.620 |

RD = root diameter, SRL = specific root length, RTD = root tissue density, RN = root nitrogen concentration.

A total of 101 studies were included in this article for data extraction:

